# Chitosan-coated carboxylic acids show antimicrobial activity against antibiotic-resistant Gram-negative and positive pathogens

**DOI:** 10.1101/2022.09.02.506354

**Authors:** Tristan Cogan, Lynn James-Meyer

## Abstract

Antibiotic resistance in bacteria is suggested to be the greatest risk to human health, but new agents are not being brought to market as the rapid evolution of resistance to them means that drug development costs cannot be recouped. Fatty acids have been proposed as a new generation of antibiotics, but toxicity and poor absorption has meant that their use has been impractical in the past. Chitosan has been used to encapsulate other agents as nanoparticles, but has not been used with fatty acids. Here we show that chitosan can be modified to direct fatty acids towards Gram-positive or negative bacteria so that they exert antimicrobial effects. We show that fatty acids work as effective antibiotics in vitro and in vivo, with activity against extremely drug resistant pathogens. Bacteria exposed to them do not develop resistance to these agents, and they are not toxic to mammalian cells. Activity was seen against salmonellosis and *C. difficile* infection in animal models. Our results demonstrate that fatty acids formulated as chitosan nanoparticles are effective antibiotics, and can be used for a long period of time without resistance developing. This suggests that the usage of fatty acids coated in this manner could be sold in sufficient quantities to recoup its development costs, overcoming this barrier. These agents would form a new class of antibiotics, with the novel property of lack of bacterial resistance.

## Introduction

The World Health Organisation has recognised a number of antibiotic-resistant pathogens as posing the greatest threat to human health. It further concluded that mortality and morbidity from resistant infections is on the rise globally, the clinical anti-bacterial pipeline remains insufficient and the outlook remains bleak (WHO 2019). A recent Wellcome Trust report concludes that it is ’no exaggeration to say that antibiotics are the foundations of modern medicine. But these foundations are crumbling’ (Wellcome Trust, 2020). This is exacerbated by economic disincentives. It typically takes 10-15 years and over $1 Billion to produce a new antibiotic.

The fact that bacteria can develop resistance within months of a new antibiotic being deployed has acted as a powerful disincentive for antibiotic development, with only two new classes being developed in the last 40 years (Coates, Halls and Hu, 2011). In the laboratory this has been illustrated to occur in a matter of days following the exposure of non-resistant bacteria to an agent in agar, resulting in the evolution of stronger resistance over time (Baym *et al.,* 2016). It is clear that new classes of antibiotic with novel targets need to be developed, but also that these need to present a barrier to the development of antibiotic resistance.

The majority of the bacteria needing new antibiotics to be developed as a critical or high priority are Gram-negative – the ESKAPE pathogens – but a minority of Gram-positives such as vancomycin-resistant *Enterococcus faecium* are also high priority.

Carboxylic acids and their derivatives are known to have weak antimicrobial activity against Gram-positive bacteria in their native state (Casillas-Vargas *et al*., 2021), with activity against Gram-negative organisms only seen at very high concentrations (Kabara *et al.,* 1972), which make them impractical for use as a therapeutic agent. Alkyl amine and amide derivatives of fatty acids (with nitrogen atoms in or attached to the hydrocarbon chain) have improved activity, with some showing weak activity against Gram-negatives, but have chronic effects making them unsuitable for use as a human chemotherapeutic agent (Anonymous, 1999).

Casillas-Vargas *et al.* (2021) suggested that fatty acids could form the basis of a new generation of antimicrobial agents due to their ability to act directly on bacteria and to synergise with existing antibiotics. Lam *et al.* (2016) had previously shown that a number of fatty acids could be formulated as phospholipid liposomes, and that these showed antibacterial activity, but that absorption into eukaryotic membranes and cytotoxic activity was a problem with extended contact, meaning that such preparations were likely to be limited to topical usage.

Bacteria have a long history of exposure to carboxylic acids, with chain lengths of up to C18 being present in milk, animal and plant tissues. Despite millenia of exposure to inhibitory or subinhibitory concentrations, carboxylic acids at high concentrations are still reliable food preservation agents (Vazquez *et al.,* 2011), with no indication that resistance has ever developed.

Carboxylic acids act on the membrane of bacteria, disrupting the function of the membrane and altering fluidity (Parsons *et al.,* 2012) and altering cell signalling pathways (Ibarguren *et al.,* 2014). It is also suggested that production of reactive oxygen species, DNA/RNA/protein synthesis inhibition and metabolic inhibition could also be mechanisms of action (Casillas-Vargas *et al.,* 2021; Xu *et al*., 2015; Zhang *et al*., 2017).

These substances are readily absorbed into the skin and mucous membranes. This property is desirable in a topical agent, but limits their efficacy in internal usage, where rapid absorbance and turnover by eukaryotic cells means that they cannot reach a distal site of action.

Nanoparticle encapsulation has been used in recent years to protect drugs from off-target absorbance, and has been used to deliver higher concentrations of antibiotics to bacteria than would be possible using the agents in their native state (Leid *et al*., 2012). This has been shown to be effective in overcoming low level resistance to agents in MRSA (Scolari *et al.,* 2020) and in *Mycobacterium* (Aboutaleb *et al.,* 2012).

Here we show that chitosan nanoparticle coating of C12 fatty acids renders them active against Gram-positive bacteria at lower concentration than free fatty acids. We show that chemical decoration of the chitosan enables these fatty acids extend their spectrum of activity to encompass the Gram-negative ESKAPE pathogens, with activity seen against highly antibiotic resistant bacteria.

Finally, we demonstrate that these compounds are non-toxic and that no resistance develops on long term exposure of bacteria to the compounds either in broth using the spatiotemporal evolution method of Baym *et al.* (2016).

We suggest that these coated carboxylic acids represent a usable new class of antibiotics with no likelihood of rapid bacterial resistance developing.

## Materials and Methods

### Synthesis of nanoparticles

All suspensions were made in a stock buffer containing 2mM sodium acetate and 98 mM acetic acid. Lecithin (22 g/l) was added to this buffer, mixed and sonicated for 1 h in a sonicating water bath so that it was totally suspended. Separately, chitosan (Sigma 448869; 15 g/l) was prepared in the same manner. Test compounds (table 1) were added to the lecithin suspension at 10% w/v and again sonicated for 1 h in a water bath. An equal volume of chitosan suspension was added to this and again sonicated. In other subsequent experiments glycol chitosan was substituted for chitosan. After sonication compounds were diluted with water up to 50% v/v in order to make a filterable suspension and passed through a 0.22 uM filter.

**Table 1.** Test compounds used with chitosan and/or glycol chitosan.

| Test compound | With chitosan | With glycol chitosan |
| --- | --- | --- |
| N-lauroyl-D-sphingosine | X |  |
| Isopropyl laurate | X |  |
| 12-(7-nitrobenzofuran-4-ylamino)dodecanoic acid | X |  |
| 4-nitrophenyl dodecanoate | X |  |
| Sucrose monolaurate | X | X |
| 1-lauroyl-rac-glycerol | X |  |
| 3-oxo-N-(2-oxocyclohexyl) dodecanamide | X |  |
| 12-aminododecanoic acid | X | X |
| 12-amino 1-dodecanoic acid methyl ester HCl | X | X |
| Pentyl laurate | X | X |
| Lauric acid | X | X |
| Ethyl dodecanoate | X | X |
| Hexyl laurate | X | X |
| Butyl laurate | X | X |
| Benzyl laurate | X | X |
| Isoamyl laurate | X | X |
| Monolaurin | X | X |
| Methyl laurate | X | X |
| Octanoic acid (positive control) | X | X |
| Decanoic acid | X | X |
| Water (negative control) | X | X |

### Bacteria used and AMR test method

Bacteria used are listed in table 2. Three strains of each species were used, sourced from laboratory collections. MICs were performed in triplicate using the EUCAST broth microdilution method using the compounds listed in table 1.

**Table 2.**
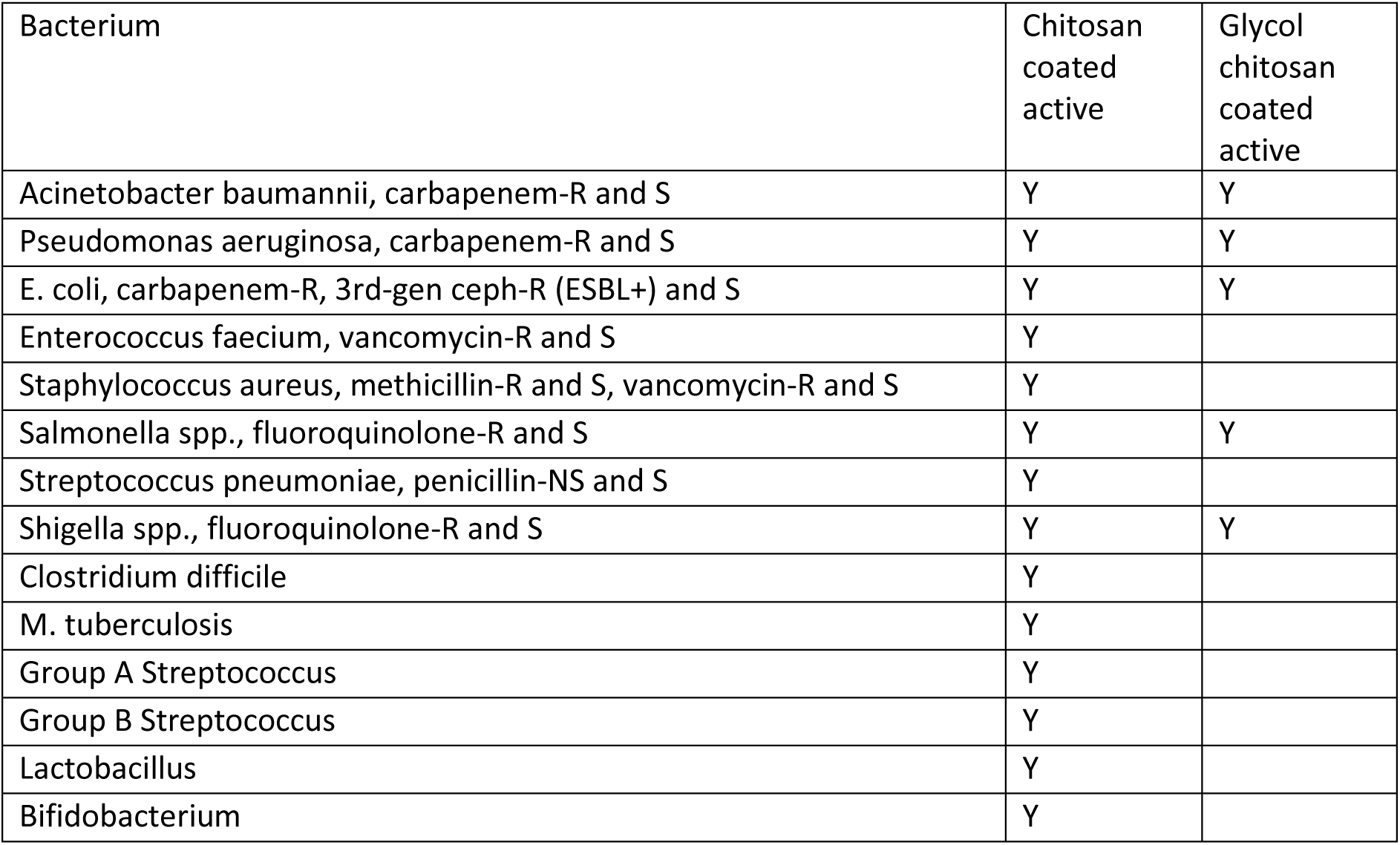
Bacteria tested.

| Bacterium | Chitosan coated active | Glycol chitosan coated active |
| --- | --- | --- |
| Acinetobacter baumannii, carbapenem-R and S | Y | Y |
| Pseudomonas aeruginosa, carbapenem-R and S | Y | Y |
| E. coli, carbapenem-R, 3rd-gen ceph-R (ESBL+) and S | Y | Y |
| Enterococcus faecium, vancomycin-R and S | Y |  |
| Staphylococcus aureus, methicillin-R and S, vancomycin-R and S | Y |  |
| Salmonella spp., fluoroquinolone-R and S | Y | Y |
| Streptococcus pneumoniae, penicillin-NS and S | Y |  |
| Shigella spp., fluoroquinolone-R and S | Y | Y |
| Clostridium difficile | Y |  |
| M. tuberculosis | Y |  |
| Group A Streptococcus | Y |  |
| Group B Streptococcus | Y |  |
| Lactobacillus | Y |  |
| Bifidobacterium | Y |  |

### Temporal analysis of development of antimicrobial resistance

Following *in vitro* assays, ethyl dodecanoate in chitosan (PY-1) and 12-amino-1-dodecanoic acid methyl ester in glycol chitosan (PY-3) were taken forward for further testing.

*Staphylococcus aureus* was assessed for the development of resistance to PY-1 by serial exposure of the bacterium to half inhibitory concentrations of chitosan-coated nanoparticulate suspensions of this compound. The bacterium and test compound were added to Mueller Hinton broth and incubated at 37°C for 7 d before subculture. At each subculture, MIC was checked by broth microdilution as above. This was continued for 6 months.

Development of resistance to PY-3 was determined using the MEGA plate assay of Baym *et al.* (2016). Concentrations of 2, 20 and 200 x the MIC were used in the plate. The plate was incubated at 30°C for 14 days.

### Toxicity

Toxicity was assessed by adding 10x MIC amounts, and tenfold dilutions of these, of compounds in table 1 to confluent monolayers of Caco-2 cells. These were incubated in a humidified CO2 atmosphere for 24 hours then cell death assessed using an LDH assay.

### In vivo activity of PY-1 against *Clostridium difficile*

All animal experiments were reviewed and approved by local ethical review committees. Hamster experiments were carried out under a Home Office licence. PY-1 (0.5g active/kg bodyweight) was trialled as a prophylactic measure to prevent *Clostridium difficile* colitis in hamsters, against a group of animals given no treatment. Hamsters were gavaged with PY-1 twice daily for one day prior to dosage with *C. difficile* followed by dosage with clindamycin to induce colitis. PY-1 treatment was then continued twice daily to assess its ability to prevent the occurrence of colitis by *C. difficile.* Evaluation of physical parameters in animals was used as a primary measure of efficacy; stool samples were also taken daily for *C. difficile* toxin analysis using a commercial SNAP test. Animals were terminated when clinical signs of disease were seen or at 10 days post infection.

### In vivo activity of PY-3 against *Salmonella*

Chickens - PY-3 (1% in water) was fed to 7-day-old Ross chicks that had been infected with *Salmonella enterica* serovar Enteritidis strain LA5 (10^4^ cfu/g feed) at day old. Birds were kept in two groups of 12 with one being dosed with PY-3 and one with water. Housing was as per breed manual recommendations. At 10 days old birds were euthanased and spleen and caecum analysed for *Salmonella.* Samples were homogenised and examined for *Salmonella* by enrichment in 9 volumes of selenite cystine broth for 18 hrs at 41°C. A 10 microlitre volume of the enriched broth was subsequently streaked on XLD agar.

Mice – Six-week-old female C57BL/6 mice were randomly allocated to experimental groups and allowed to acclimatise for a week. On Day 0, animals were fasted for 4 hr (food and water) before per os (p.o.) treatment with 20 mg of streptomycin. 72 h after streptomycin treatment, mice were infected with *Salmonella* (10^4^ cfu in PBS p.o.); one group was provided with food and water (untreated) ad libitum and the remaining group received food and water plus 1% PY-3) ad libitum. 72 h post infection, mice were sacrificed and infection determined from faecal pellets using methods as for chicks above.

All studies were carried out under licence and approved by the relevant local ethics committee.

## RESULTS

A number of the fatty acid derivatives tested showed activity against Gram-positive pathogens, but not Gram-negative pathogens or the commensal bacteria *Bifidobacterium* or *Lactobacillus* (Table 3). Antibiotic sensitive (Table 3) and resistant (Table 4) organisms shows identical MICs. Among the fatty acids with 12 carbon backbones, sucrose monolaurate showed activity at the highest dilution, but suspensions showed a high level of viscosity (data not shown), so this compound was not taken forward for further work. Methyl laurate showed activity against the Gram-positive agents tested, including *Mycobacterium tuberculosis*, so this formulation was taken forward for further study.

**Table 3.**
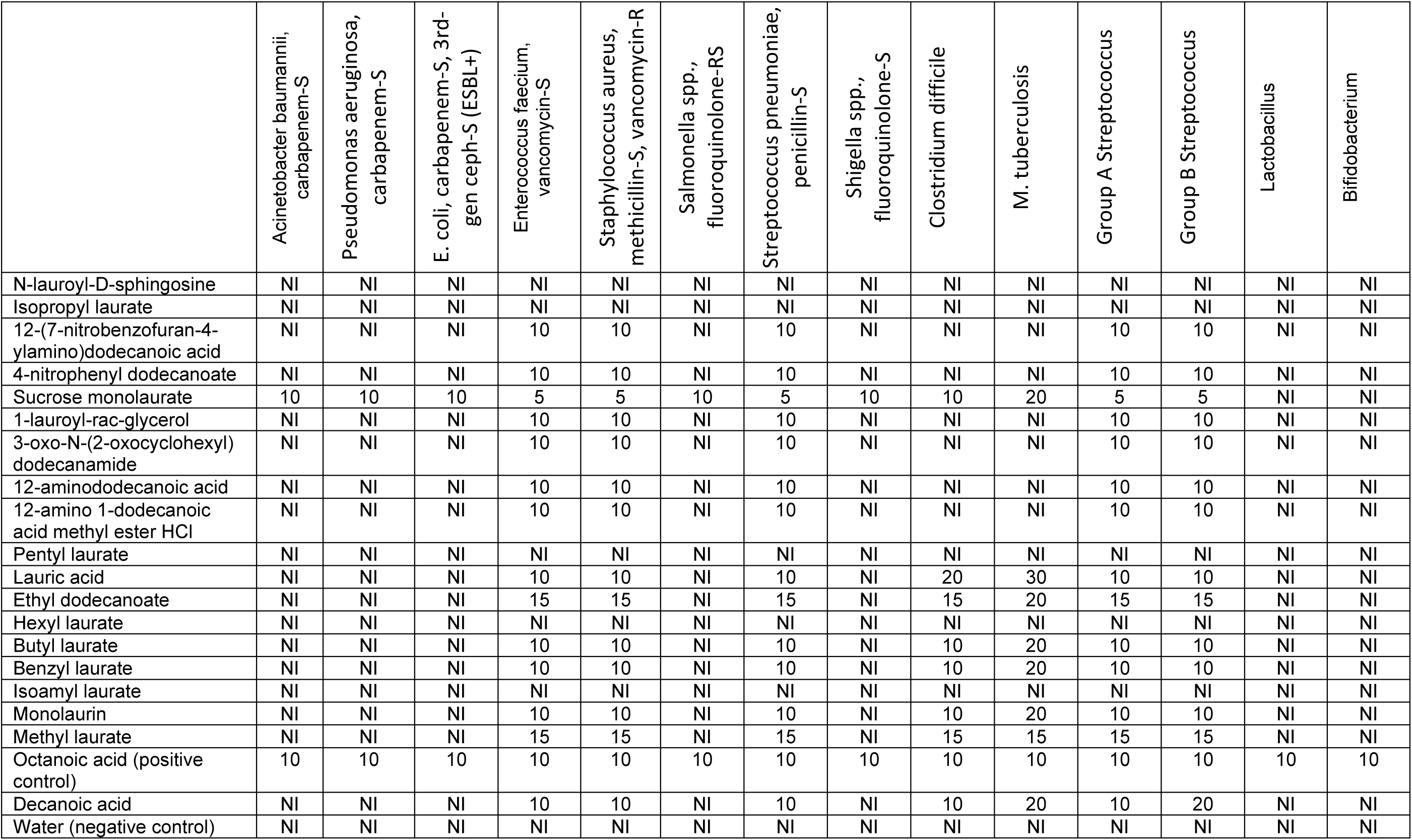
MICs of chitosan-coated fatty acids against test pathogens in µg/ml. NI – no inhibition.

**Table 4.** MICs of chitosan-coated fatty acids against resistant test pathogens in µg/ml. NI – no inhibition.

|  | Acinetobacter baumannii,<br>carbapenem-R | Pseudomonas aeruginosa,<br>carbapenem-R | E. coli, carbapenem-R, 3rd-<br>gen ceph-R (ESBL+) | Enterococcus faecium,<br>vancomycin-R | Staphylococcus aureus,<br>methicillin-R, vancomycin-R | Salmonella spp.,<br>fluoroquinolone-R | Streptococcus<br>pneumoniae, penicillin-NS | Shigella spp.,<br>fluoroquinolone-R |
| --- | --- | --- | --- | --- | --- | --- | --- | --- |
| N-lauroyl-D-sphingosine | NI | NI | NI | NI | NI | NI | NI | NI |
| Isopropyl laurate | NI | NI | NI | NI | NI | NI | NI | NI |
| 12-(7-nitrobenzofuran-4-ylamino)dodecanoic acid | NI | NI | NI | 10 | 10 | NI | 10 | NI |
| 4-nitrophenyl dodecanoate | NI | NI | NI | 10 | 10 | NI | 10 | NI |
| Sucrose monolaurate | 10 | 10 | 10 | 5 | 5 | 10 | 5 | 10 |
| 1-lauroyl-rac-glycerol | NI | NI | NI | 10 | 10 | NI | 10 | NI |
| 3-oxo-N-(2-oxocyclohexyl)<br>dodecanamide | NI | NI | NI | 10 | 10 | NI | 10 | NI |
| 12-aminododecanoic acid | NI | NI | NI | 10 | 10 | NI | 10 | NI |
| 12-amino 1-dodecanoic<br>acid methyl ester HCl | NI | NI | NI | 10 | 10 | NI | 10 | NI |
| Pentyl laurate | NI | NI | NI | NI | NI | NI | NI | NI |
| Lauric acid | NI | NI | NI | 10 | 10 | NI | 10 | NI |
| Ethyl dodecanoate | NI | NI | NI | 15 | 15 | NI | 15 | NI |
| Hexyl laurate | NI | NI | NI | NI | NI | NI | NI | NI |
| Butyl laurate | NI | NI | NI | 10 | 10 | NI | 10 | NI |
| Benzyl laurate | NI | NI | NI | 10 | 10 | NI | 10 | NI |
| Isoamyl laurate | NI | NI | NI | NI | NI | NI | NI | NI |
| Monolaurin | NI | NI | NI | 10 | 10 | NI | 10 | NI |
| Methyl laurate | NI | NI | NI | 15 | 15 | NI | 15 | NI |
| Octanoic acid (positive<br>control) | 10 | 10 | 10 | 10 | 10 | 10 | 10 | 10 |
| Decanoic acid | NI | NI | NI | 10 | 10 | NI | 10 | NI |
| Water (negative control) | NI | NI | NI | NI | NI | NI | NI | NI |

Gram-negative pathogens were not inhibited by any of the fatty acids in chitosan that were tested, except for octanoic acid. This was used as a positive control as it is known to have antibacterial activity but is indiscriminate in its activity.

A number of compounds were taken forward in a formulation utilising glycol chitosan coated nanoparticles (Tables 5 and 6). Three of these compounds showed activity against Gram negatives, with no difference seen in MICs against antibiotic sensitive and resistant pathogens. 12-aminododecanoic acid was taken forward for further study as it had low viscosity and activity at high dilution.

**Table 5.** MICs of glycol chitosan-coated fatty acids against test pathogens in µg/ml. NI – no inhibition.

|  | Acinetobacter baumannii,<br>carbapenem-S | Pseudomonas aeruginosa,<br>carbapenem-S | E. coli, carbapenem-S, 3rd-<br>gen ceph-S (ESBL+) | Salmonella spp.,<br>fluoroquinolone-RS | Streptococcus pneumoniae,<br>penicillin-S | Shigella spp.,<br>fluoroquinolone-S | Lactobacillus | Bifidobacterium |
| --- | --- | --- | --- | --- | --- | --- | --- | --- |
| Sucrose monolaurate | 1 | 1 | 1 | 1 | 1 | 1 | NI | NI |
| 12-aminododecanoic acid | 1 | 1 | 1 | 1 | 1 | 1 | NI | NI |
| 12-amino 1-dodecanoic<br>acid methyl ester HCl | 5 | 5 | 5 | 5 | 5 | 5 | NI | NI |
| Pentyl laurate | NI | NI | NI | NI | NI | NI | NI | NI |
| Lauric acid | NI | NI | NI | NI | NI | NI | NI | NI |
| Ethyl dodecanoate | NI | NI | NI | NI | NI | NI | NI | NI |
| Hexyl laurate | NI | NI | NI | NI | NI | NI | NI | NI |
| Butyl laurate | NI | NI | NI | NI | NI | NI | NI | NI |
| Benzyl laurate | NI | NI | NI | NI | NI | NI | NI | NI |
| Isoamyl laurate | NI | NI | NI | NI | NI | NI | NI | NI |
| Monolaurin | NI | NI | NI | NI | NI | NI | NI | NI |
| Methyl laurate | NI | NI | NI | NI | NI | NI | NI | NI |
| Octanoic acid | NI | NI | NI | NI | NI | NI | NI | NI |
| Decanoic acid | NI | NI | NI | NI | NI | NI | NI | NI |
| Water (negative control) | NI | NI | NI | NI | NI | NI | NI | NI |

**Table 6.** MICs of glycol chitosan-coated fatty acids against resistant test pathogens in µg/ml. NI – no inhibition.

|  | Acinetobacter baumannii,<br>carbapenem-R | Pseudomonas aeruginosa,<br>carbapenem-R | E. coli, carbapenem-R, 3rd-<br>gen ceph-R (ESBL+) | Enterococcus faecium,<br>vancomycin-R | Salmonella spp.,<br>fluoroquinolone-R | Shigella spp.,<br>fluoroquinolone-R |
| --- | --- | --- | --- | --- | --- | --- |
| Sucrose monolaurate | 1 | 1 | 1 | 1 | 1 | 1 |
| 12-aminododecanoic acid | 1 | 1 | 1 | 1 | 1 | 1 |
| 12-amino 1-dodecanoic<br>acid methyl ester HCl | 5 | 5 | 5 | 5 | 5 | 5 |
| Pentyl laurate | NI | NI | NI | NI | NI | NI |
| Lauric acid | NI | NI | NI | NI | NI | NI |
| Ethyl dodecanoate | NI | NI | NI | NI | NI | NI |
| Hexyl laurate | NI | NI | NI | NI | NI | NI |
| Butyl laurate | NI | NI | NI | NI | NI | NI |
| Benzyl laurate | NI | NI | NI | NI | NI | NI |
| Isoamyl laurate | NI | NI | NI | NI | NI | NI |
| Monolaurin | NI | NI | NI | NI | NI | NI |
| Methyl laurate | NI | NI | NI | NI | NI | NI |
| Octanoic acid | NI | NI | NI | NI | NI | NI |
| Decanoic acid | NI | NI | NI | NI | NI | NI |
| Water (negative control) | NI | NI | NI | NI | NI | NI |

Antibacterial activity appeared to be primarily mediated by the agents themselves, with the form of chitosan used acting as a vehicle to deliver these to the bacteria, and selecting whether Gram positives or Gram negatives were targeted.

Following *in vitro* assays, ethyl dodecanoate in chitosan (PY-1) and 12-amino-1-dodecanoic acid methyl ester in glycol chitosan (PY-3) were taken forward for further testing.

Over a 6 month period of exposure to PY-1, no change was seen in the MIC of this compound for *Staphylococcus*, indicating an inability of this bacterium to evolve resistance to PY-1; PY-3 was not tested in this assay. The same results were seen with MEGA plates, on which *E. coli* were not able to evolve resistance to PY-3, with no bacteria growing beyond the agar zone containing no antibacterial agent (data not shown).

No cytotoxicity was seen on LDH assay of Caco-2 cells exposed to PY-1 or PY-3 (data not shown).

PY-1 (0.5g active/kg bodyweight) was trialled as a prophylactic measure to prevent *C. difficile* colitis in hamsters against a group of animals given no treatment. There was no effect of treatment on the number of animals needing to be euthanased; these fell into two groups: those that were moribund, and those showing ‘wet tail’ – an indication of *C. diff.* toxin-mediated diarrhoea. 5/10 animals in the control group had wet tail, compared to 2/10 in the treated group. This difference in diarrhoea was supported by *C. difficile* toxin being detected in 6/10 control animals beyond day 7, against 1/10 PY-1 treated animals (Fishers exact test P=0.028, significant difference between groups; Figure 1).

**Figure 1.**
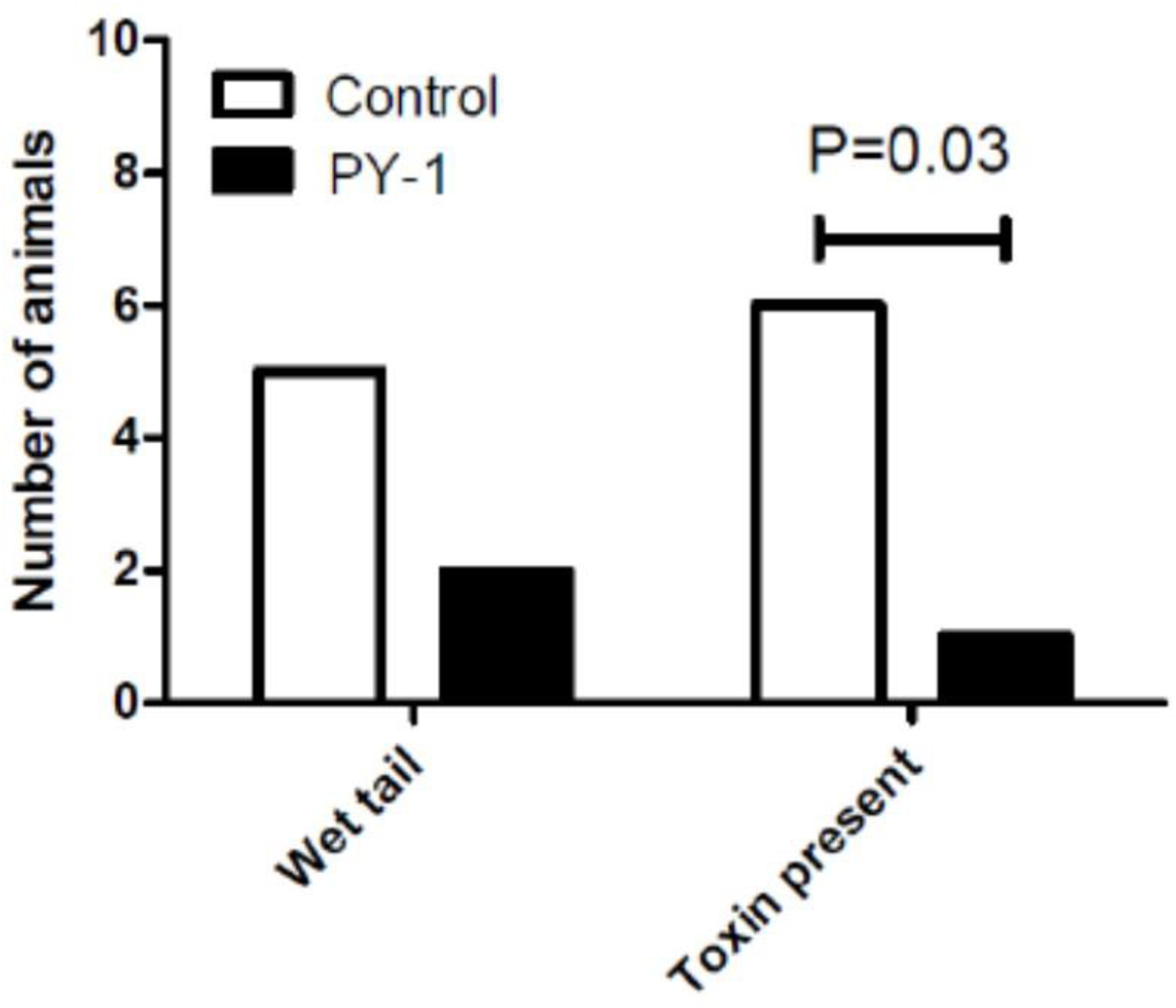
Occurrence of wet tail and toxin secretion in hamsters infected with *C. difficile* and treated with PY-1 or control. N=15 animals per group.

PY-3 showed effectiveness in removing *Salmonella* colonisation in caecum in the chick model (Fishers exact test P=0.04, significant difference between groups; Figure 2). No colonisation of the spleen was observed in either group, suggesting a lack of systemic invasion. Similar effectiveness was seen in the mouse (Fishers exact test P=0.0498, significant difference between groups; Figure 3).

**Figure 2.**
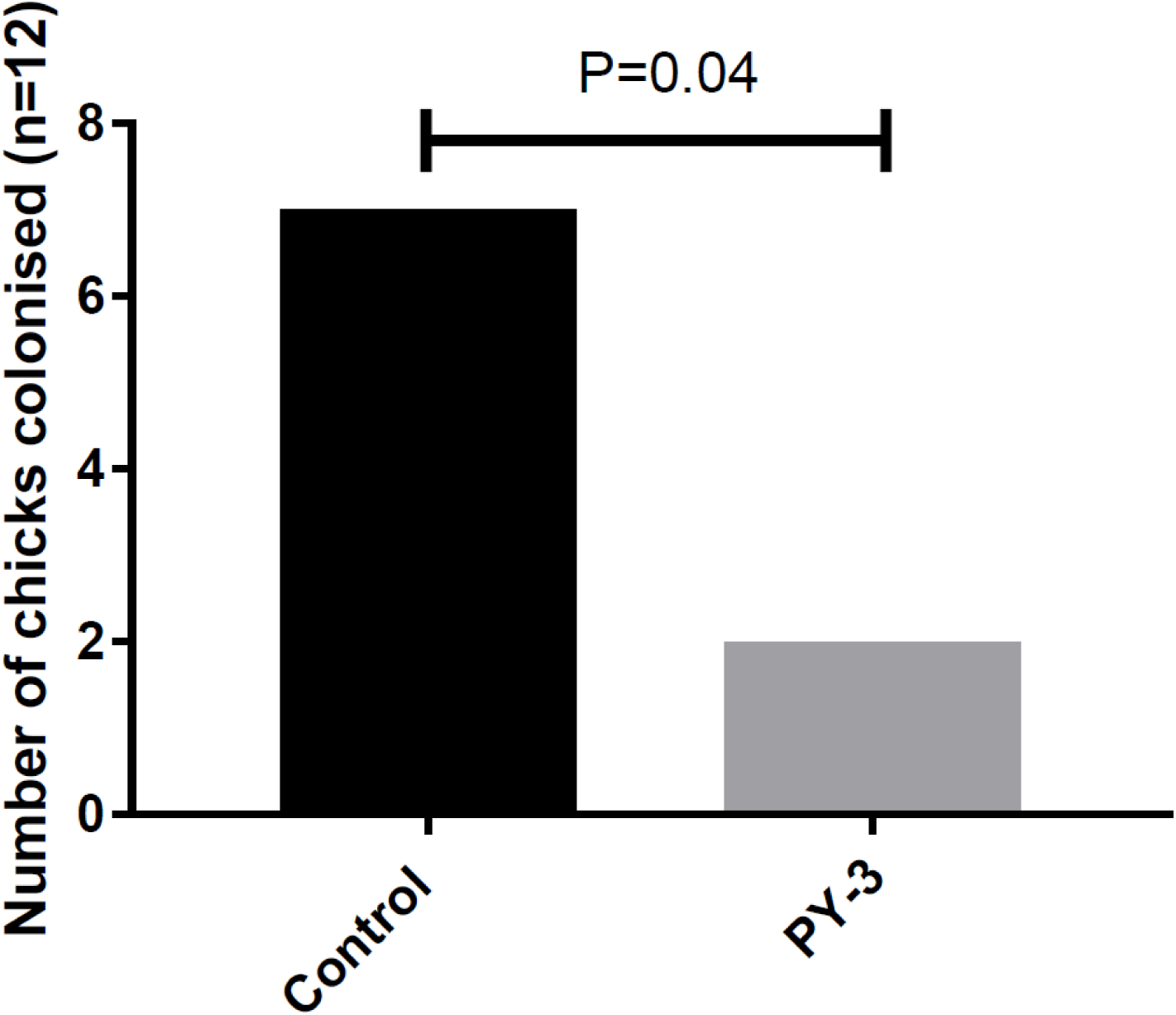
Removal of *Salmonella* colonisation of the caecum in a chick model after 3 d dosing with PY-3. N=12 animals per group.

**Figure 3.**
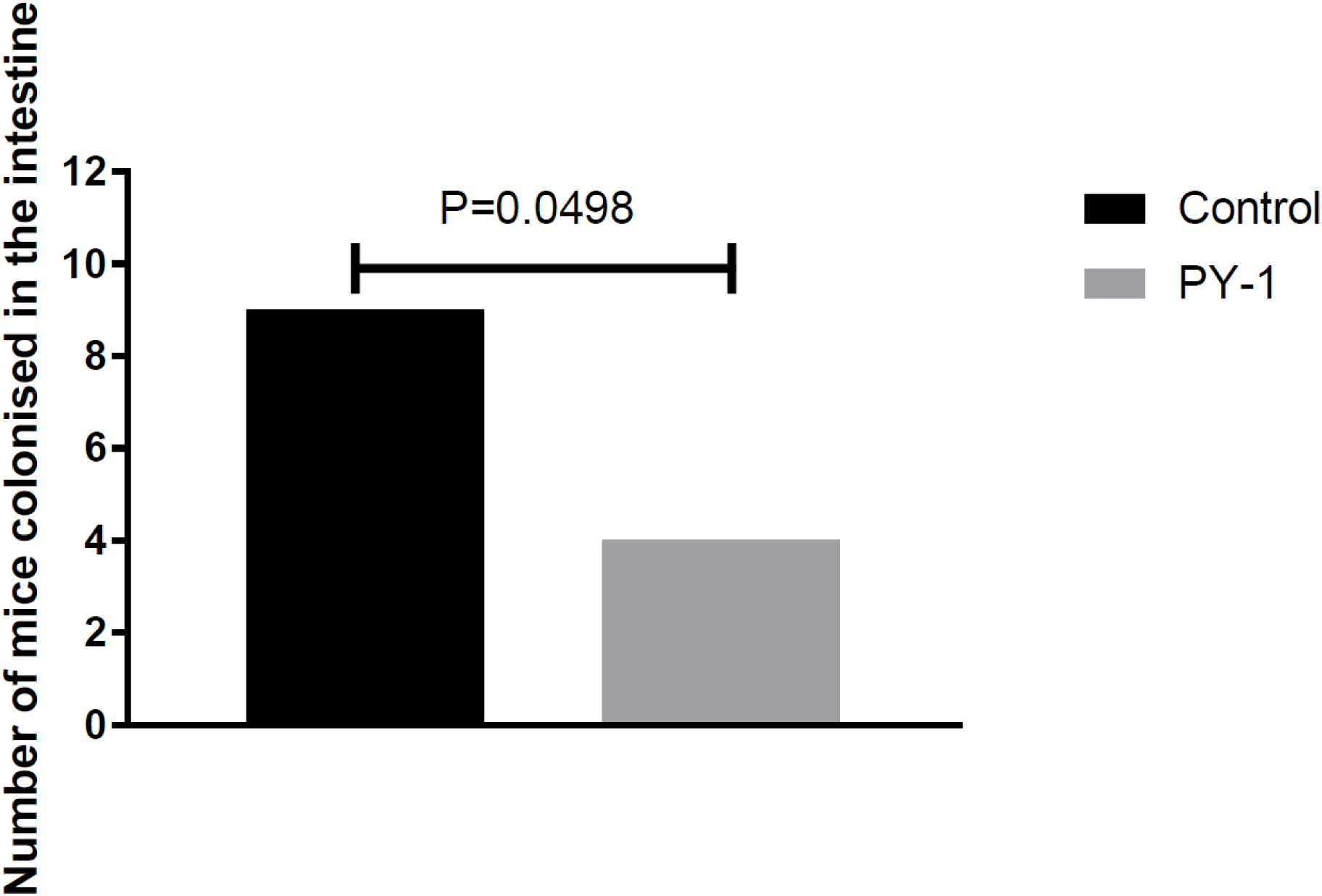
Removal of *Salmonella* colonisation of the intestine in streptomycin-treated mice after 3 d dosing with PY-3. N=12 animals per group.

No animals dosed with either PY-1 or 3 showed any signs of ill health or negative reactions to dosing.

## DISCUSSION

It has recently been suggested that fatty acids could be utilised as antibiotics (Casillas-Vargas *et al.,* (2021). The high concentrations of native fatty acids needed to exert this effect (Kabara *et al.,* 1972) and issues with toxicity (Anonymous, 1999, Lam *et al.,* 2016) mean that any uses other than topical are problematic, limiting their usage against the highest priority antibiotic resistant ESKAPE pathogens. This work has shown that nanoparticulate encapsulation of fatty acids by chitosan and its derivatives enables these compounds to be used as antimicrobials at pharmacologically-relevant concentrations.

Chitosan coating was found to be effective at targeting the active compounds tested to Gram-positive bacteria, but not Gram negatives (Tables 3 and 4). It has been shown before that Gram-negative bacteria are less susceptible to fatty acid insertion that Gram-negatives as the outer membrane of these bacteria is protective (Miller *et al.,* 1977). The conjugation of glycol to chitosan enabled the targeting of Gram negative pathogens (Tables 5 and 6). The key mediator of activity was found to be the active fatty acid, rather than the encapsulating compound, as many fatty acids were inactive.

In order to achieve widespread usage it is important that an antibiotic should not encourage the development of resistance in bacteria, and that it should be active against bacteria exhibiting existing mechanism of resistance to other classes of antibiotic. Existing MDR and sensitive bacteria showed the same MICs to the test compounds (Tables 3 vs. 4 and 5 vs. 6), and existing MDR bacteria did not show resistance to either chitosan or glycol chitosan coated fatty acids where there was any activity (Tables 4 and 6), indicating that currently circulating mechanisms of antibiotic resistance do not cause resistance to these agents.

Furthermore, the fact that fatty acids target multiple cellular processes and pathways, including energy generation, nutrient uptake and induction of autolysis (reviewed by Desbois and Smith, 2010) limits the ability of bacteria to simultaneously evolve resistance across a number of targets.

Extended incubation of bacteria with PY-1 and 3 showed no evolved resistance developing (data not shown), while the MEGA plate assay of Baym *et al.* (2016) also did not show that resistance could develop.

Fatty acids are thought to intercalate into the bacterial membrane (Casillas-Vargas *et al.,* 2021, Miller *et al.,* 1977), where they would impair membrane fluidity, function and toxin secretion. This is supported by our data in Figure 1, where toxin secretion by *C. difficile* was inhibited by PY-1 dosing, hinting at a mechanism of action. Branched fatty acids disorder membrane packing. *Lactobacillus* and *Bifidobacterium*, having evolved to grow in milk which is high in fatty acids possess high levels of enzymes which can alter branched chain fatty acids in their membranes (Veerkamp, 1971), as do mammalian cells. Pathogenic bacteria do not possess these pathways and so are susceptible. To evolve resistance, pathogens would have to alter their membrane composition and acquire enzyme pathways that could interact with and alter branched chain fatty acids within their membrane, thus altering their growth temperature and becoming ineffective pathogens. For this reason, we believe that resistance to PY-1 and 3 cannot develop in pathogens, meaning that they could have widespread use as antimicrobials without the worry that resistance would develop.

The fact that mammalian cells can metabolise these compounds is also the key to their safety, as we saw here there was no toxicity. It does, though, mean that more efficient targeting of bacteria will rely on encapsulation of these agents, or antibody-mediated targeting of pathogens. This is the subject of further research and development.

Evidence of *in vivo* efficacy was seen in small-group experiments looking at the effect of PY-1 on *C. difficile* in mice and PY-3 against *Salmonella* in chicks and mice. No adverse effects were seen in these experiments, indicating that orally-dosed fatty acids could be employed as a pharmaceutical treatment for disease. Larger studies are clearly necessary, though.

In summary, we have shown that chitosan encapsulation can enable fatty acids to be used as antimicrobials that will efficiently target the highest priority ESKAPE pathogens. The results of this work indicate that fatty acids, suitably coated, have huge potential as safe antimicrobials to which resistance cannot develop.

